# ScPectral: Spectrally Clustering Hypergraph Representations of Transcription Networks to Identify Developmental Pathways

**DOI:** 10.1101/2024.12.19.629530

**Authors:** Dennis Bersenev

**Affiliations:** The Systems Biology Institute

## Abstract

Transcription Networks, otherwise known as Gene Regulatory Networks (GRNs), are models of biological systems centred on Transcription Factor (TF) interactions. These models equip experimentalists with a powerful computational tool to predict the effects of different genetic perturbations. GRNs are canonically modelled using a digraph, wherein the arcs indicate activation or repression between each pair of nodes to represent the relationships among the TFs. However, gene regulation is accomplished by groups of TFs working in concert, a biological reality the pairwise model neglects. In addition to the paucity of GRN representations incorporating this known TF biology, a persisting challenge to inference of the networks themselves is in accounting for the latent dynamics of gene interactions. In considering this second point, the advent of single-cell RNA sequencing technologies, provides the high resolution data needed to begin effectively inferring temporally-aware models. Despite this, utilisation of temporally-aware statistical metrics to do so has been limited. In addressing these shortcomings to GRN inference, scPectral is introduced as a method to infer a robust dynamic representation of a common GRN motif, the cascade, in the form of a hypergraph. ScPectral is applied to the identification of developmental pathways for known processes to validate its efficacy. Given scPectral’s modest success in finding key constituents of developmental pathways, and its ability to do so in a manner requiring no input or annotation of known biology, through further improvement it may develop to become a technique able to aid experimentalists exploring novel development processes. ScPectral is made available at: https://github.com/Dennis-Bersenev/scPectral.

## Introduction

In the field of Systems Biology, the cell is viewed as an information processing unit that is continuously monitoring its environment, and determining which proteins are needed in what amounts (Alon 2020). Of particular import are special proteins called transcription factors, TFs, which regulate gene expression and are thereby used to represent the state of the cell (Alon 2020). Gene Regulatory Networks (GRNs) model the interactions among transcription factors and their target genes, in effect they describe the dynamics of the cell (Alon 2020). Inference of GRNs therefore, can potentially provide significant insights into the mechanisms underlying cellular processes (Stumpf 2021). Thus far, however, the inference of GRNs, and deriving meaningful biology from them remains an unsolved challenge (Stumpf 2021). The principal problem with inferring gene regulatory relationships is that the networks are intrinsically dynamic, while the sequencing data used to infer them is static (Stumpf 2021). Currently, the technique most often applied in addressing this issue is Trajectory Inference, TI (Luecken and Theis 2019). Unfortunately, TI is a technique that largely fails in its objective of providing a pseudotemporal ordering as effective as true time series data in reconstructing GRNs (Qiu et al. 2020). In addition to the limitations of TI techniques, GRNs are most commonly modelled as pairwise networks; however, the interplay among TFs is known to extend beyond pairs of individual interactions (Stumpf 2021).

Given the nature of transcription factors to work in concert across complex multi-stage processes, modelling their interactions as a pairwise graph with edges weighted by static statistics such as mutual information (Heydari et al. 2022, Qiu et al. 2020, Kim et al. 2021) is severely limiting. Such models have demonstrable shortcomings when it comes to predicting the behaviour of biological systems (Purnick and Weiss 2009). In reconciling the aforementioned challenges to modelling GRNs, the method presented: scPectral, incorporates precisely these two often neglected facts of biology. Namely, that TF interactions involve groups, not pairs, and that GRNs are dynamic entities perpetually evolving to reflect the conditions of their environment (Stumpf 2021).

ScPectral addresses these limitations by incorporating a temporally-aware information theoretic measure to build a weighted complete digraph. This digraph becomes the input to an algorithm to identify developmental cascade motifs (Figure 1) within it, the components of which form a set of hyperedges, which are ultimately clustered to identify developmental pathways. The datasets used in this analysis are of developmental processes (Hayashi et al. 2018, Marot-Lassauzaie et al. 2022). Chosen due to their well characterised biology, suitable for validating any GRN inference method. The constituent observations of these datasets are seen to represent the state of a cell as it undergoes the process of differentiation. In other words, each observation represents a different state of the cell along its developmental trajectory.

**Fig. 1.**
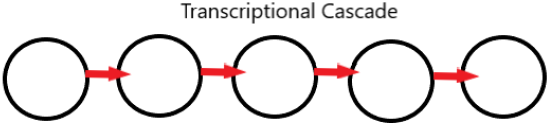
Developmental cascade

The final pathways scPectral discovers lose much of the nuance and specific interactions therein; however, this low resolution method is in large part motivated by the necessity of existing GRN inference approaches (Heydari et al. 2022) to incorporate significant a priori biological information. The limitation of these being that novel biological systems, without known pathways, cannot be discovered by such methods. ScPectral, on the other hand, requires no a priori input concerning marker genes or interactions, makes no assumptions on pseudotime ordering, and has no restrictions on either the size or number of potential pathways it discovers. ScPectral, then, is a technique which can aid experimentalists during the initial exploratory stages of investigating novel development processes.

## Methods

### Transfer Entropy

The dynamics underlying GRNs are incorporated into scPectral by using, as initial input, a collection of pairwise graphs with edges weighted by the temporally aware statistical measure of transfer entropy, TE (Thomas 2000). TE, unlike the more common information-theoretic metric of mutual information, focuses on the rate information is exchanged among the constituents of multi-component systems. The power of TE is that it can capture both dynamic and directional information (Zhang and Stumpf 2023). The pairwise graph used as input to scPectral is obtained from a recent method, locaTE (Zhang and Stumpf 2023), which models cell-state transitions as a tensor, G, of TE scores. The rank three tensor G has shape m x n x n, where m is the number of cells and n the number of genes. The index i, j, k into G tells you the transfer entropy score of j→k in the neighbourhood of cell i. These local transfer entropy scores themselves are computed according to the following equation:

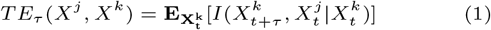

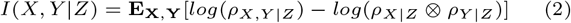

These define the transfer entropy from gene j to gene i with time lag Tau for the cell with expression matrix X (Zhang and Stumpf 2023).

### Hypergraph Extraction

ScPectral’s first step is to abstract G, as supplied by locaTE, into a collection of hyperedges and their constituent vertices to define a hypergraph. Hypergraphs are generalisations of graphs, wherein the edges (hyperedges) can connect an arbitrary number of vertices (Michoel and Nachtergaele 2012).

Formally, a hypergraph is defined as HG = (V, E), where V denotes the vertex set and E the edge set; each edge has an associated weight: *e ∈ E, w*(*e*) ∈ ℝ.

The only difference between a hypergraph and a conventional graph is that each element of the edge set E, can be a set of arbitrary cardinality; whereas, for the conventional graph each edge in E has a cardinality of two (Michoel and Nachtergaele 2012). ScPectral’s hypergraph, then, can define the interaction among a set of TFs in a more biologically accurate manner, by representing each such interaction with a single hyper edge. Each of these hyperedges being hypothesised to represent some component of a developmental process.

The constituents of each hyperedge are those vertices found along a high cost acyclic path through locaTE’s pairwise network. High cost is defined as any local TE score in *G*_*X*_ which is within some threshold of the maximum TE score for the cell X. The threshold is chosen as the highest, most restrictive, possible value while still ultimately obtaining a connected hypergraph. This edge extraction algorithm consists of seven steps:

1. Find the max TE score in locaTE’s adjacency matrix for cell X, and define a tolerance based on this score
2. Initialise each gene as a potential tail of a hyperedge set
3. For each tail, define its heads as all genes with which it has a TE score *>* tolerance
4. Merge tails with common heads into a combined edge, and remove heads that were part of that intersection from the edges involving the tails individually (Figure 2)

6. Repeatedly merge newly created edges from previous merger iterations till only a single input edge remains
7. Intersecting all heads from this step provides the set of input tails to the next iteration of the algorithm at step 3
8. Repeat 3-7 till the intersection of heads at step 7 produces no unique vertices
9. Repeat for all cells to obtain the full hyperedge set of HG

**Fig. 2.**
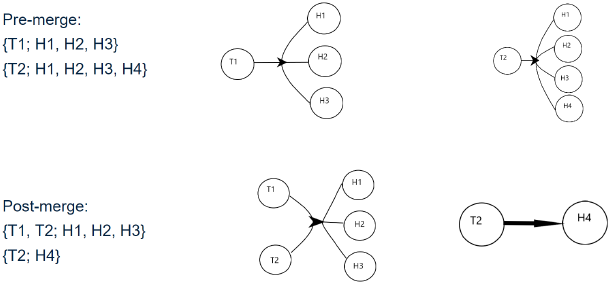
Merger graphic

Once the algorithm completes, overlapping hyperedges are those which contain vertices (genes) in common; therefore, any two hyperedges sharing a vertex are connected through that vertex, indicating they belong to the same biological pathway and each of them represent a different stage of that pathway. Connecting all such hyperedges and isolating them from others, by means of a connectivity based clustering algorithm, should then recapitulate the full pathway through time.

Spectral Clustering

With HG obtained, represented by its |*V* |*×*|*E*| incidence matrix H, spectral clustering is the connectivity based algorithm of choice used to identify the strongly connected communities within it. The implementation used was based on Ng, Jordan, and Weiss’ formulation of spectral clustering which first finds the normalised graph Laplacian, then normalises its k largest Eigenvalues to unit length, and finally performs k-means on the rows of this m x k ‘Eigenmatrix’ (Ng et al. 2001). The extension of this algorithm used for hypergraphs defines the normalised graph Laplacian as:

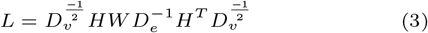

Where *D*_*v*_ and *D*_*e*_ are the diagonal vertex and edge degree matrices, respectively, and W is the diagonal matrix of hyperedge weights (Zhou and Schölkopf 2007). The spectral clustering algorithm outlined above is implemented in the HyperNetX package (Hayashi et al. 2020), which final analysis was based on. The output being a collection of clusters, each potentially defining a developmental pathway.

### Benchmarking

An analysis based on an extensive overview of the relevant biological literature, supplemented with experimental investigation in the form of perturbation studies (Heydari 2022) is ultimately needed to determine whether any of the clusters scPectral discovers are indicative of any meaningful biology. However, in using well characterised data sets with known biology and existing studies that have performed the aforementioned experimental work, a first approximation of how scPectral performs, at least in certain contrived test cases, can be done. In particular, two data sets based on stem cell differentiation were used. The first, that of murine embryonic stem cell differentiation and the second, that of murine hematopoiesis. In addition to a comparison with the existing literature on these well characterised processes, scPectral’s final results were also run through Metascape (Zhou et al. 2019) to perform enrichment analysis. Pathway enrichment analysis is a procedure to identify biological pathways that are enriched in a given list of genes (Zhou et al. 2019). Each cluster scPectral found was used as an input to Metascape’s express analysis tool, and the interpretation of each cluster as a potential biological pathway was based on these results. Through a combination of comparison to existing approaches (Hayashi et al. 2018, Zhang and Stumpf 2023, Heydari et al. 2022), and pathway enrichment through Metascape (Zhou et al. 2019) scPectral’s efficacy was determined.

## Results

### Mouse Embryonic Stem Cell Differentiation

The first dataset considered for analysis was a time-series of 456 mouse embryonic stem cells (mESCs) with 100 transcription factors, and samples collected at 0, 12, 24, 48, 72h (Hayashi et al. 2018). Specifically, this dataset captures the process of differentiation from mESCs to cells of the primitive endoderm; which in turn, under normal biological conditions, would form the linings of the respiratory and digestive tracts (Gilbert 2000). This linear differentiation trajectory was inferred by using stationaryOT (Zhang et al. 2021) to obtain the transition matrix locaTE uses to construct its pairwise network. The 456x100x100 array of interaction scores locaTE finds for the mESC dataset then becomes the input to scPectral. The hyperparameters were set to 0.75 and 5, respectively. The first, being the tolerance, indicates that any interaction score within 25% of the maximum TE for that cell is considered sufficiently meaningful to include in a pathway, this value being chosen because it was the most restrictive tolerance which still produced a connected hypergraph. While the number of clusters were selected based on the Eigengap heuristic (Luxburg 2007), seen in Figure 3.

**Fig. 3.**
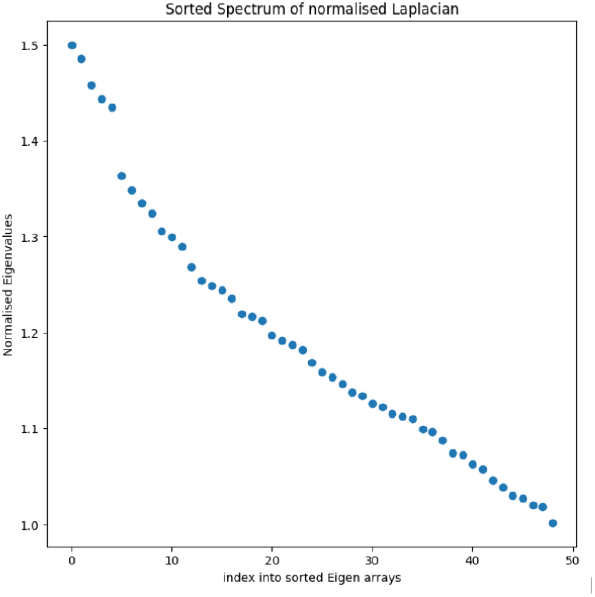
mESC sorted Laplacian Spectrum

The hypergraph was constructed according to the algorithm outlined in the methodology section, and the five resulting clusters were used to perform pathway enrichment analysis with Metscape (Zhou et al. 2019). A summary of Metascape’s results can be seen in Figure 4.

**Fig. 4.**
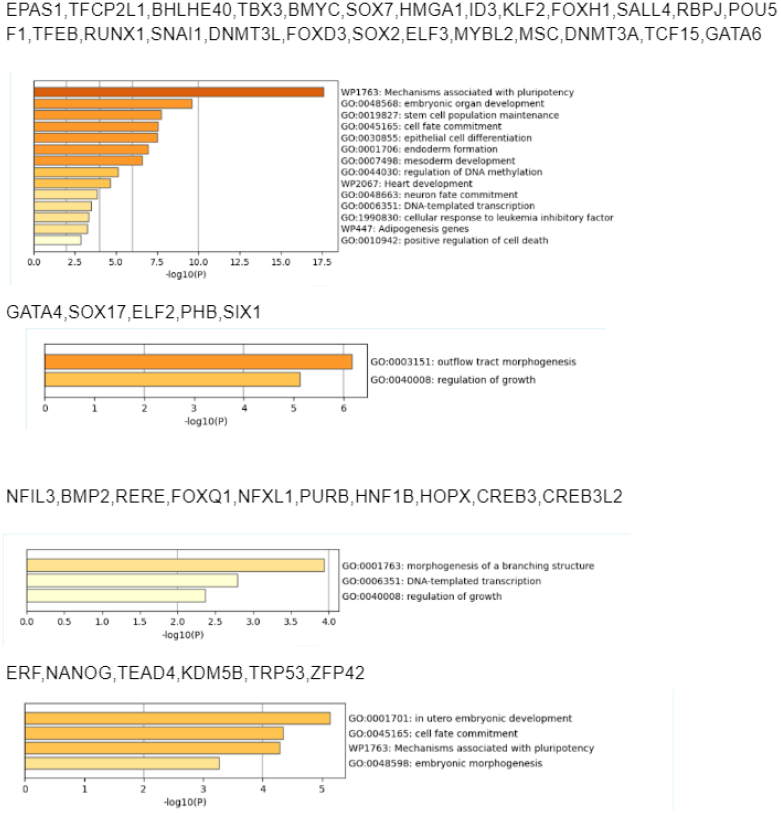
mESC Metascape

Unlike the analysis performed by Hayashi et al. or by the authors of locaTE, the goal of scPectral is not to find pathways enriched at different developmental stages; instead, it is to find pathways which span the differentiation process. In other words, scPectral finds markers of each developmental stage and includes them in a single cluster, under the hypothesis the constituents of this cluster together form a developmental cascade. Of the five potential pathways spectral clustering revealed, one was found to not be overrepresented in any known biological process, while the others were all involved in expected differentiation-related pathways (see Figure 4). Cluster 1 in particular is exemplary, as the genes it contained were overrepresented in known developmental subprocesses that include the early, mid, and late stages of development. For example, cluster 1 was enriched for mechanisms of pluripotency and stem cell population maintenance, known early markers (Hayashi et al. 2018). In addition to mid stage processes such as cell fate commitment and cell differentiation (Hayashi et al. 2018). And finally, cluster 1 genes were also significant in hallmark late stage pathways such as organ, endoderm, and mesoderm formation (Hayashi et al. 2018). The Metascape enrichment analysis therefore indicates that cluster 1 is composed of genes involved in all stages of development (Hayashi et al. 2018, Bassalert et al. 2018).

Cluster 3 is also of note, as its constituents are involved in pluripotency mechanisms, cell fate commitment, and organ development. Again, spanning the entire development process (Hayashi et al. 2018). Crucially, this cluster includes the known interaction between NANOG and ZFP42, while excluding GATA6 (Wenjing 2006). NANOG and ZFP42 are known markers along the pluripotent cell’s differentiation trajectory, and importantly NANOG and GATA6 inhibit one another while both facilitating similar developmental processes; in other words, these genes are involved in mutually exclusive processes and therefore should not appear in the same cluster (Hermitte and Chazaud 2014).

### Haematopoiesis

The second analysis investigated the extensively studied process of hematopoiesis in mice. Originally used to demonstrate the advantages of the modified RNA velocity method, *κ*-velo (Marot-Lassauzaie et al. 2022), the dataset consists of 2430 murine hematopoietic stem and progenitor cells (HSPCs), each with 50 transcription factors. The authors of locaTE used this dataset to infer the dynamics computed by *κ*-velo. That pairwise network once again became the input to scPectral. After hyper-parameter selection, again using the heuristic of a connected hypergraph with a clear Eigengap in the normalised graph Laplacian’s spectrum (Figure 5), final clustering results were obtained and used for further analysis.

**Fig. 5.**
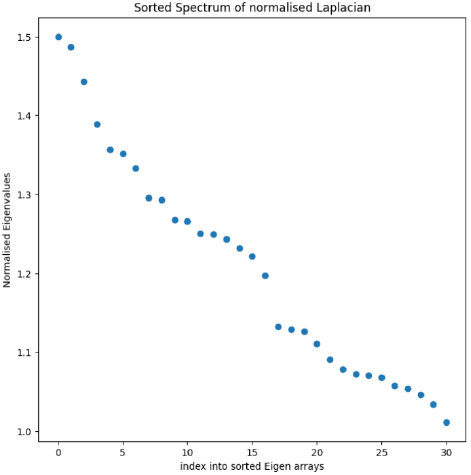
HSPC sorted Laplacian Spectrum

Haematopoiesis is an especially suitable candidate to analyse any method purporting developmental pathway identification due to the extensively studied GATA-mediated interaction network controlling Megakaryocyte–Erythroid progenitor cell fate (Doré and Crispino 2011). The key constituents of this network are the Erythroid Krüppel-like family of transcription factors (EKLF), the GATA transcription factors, and finally the myeloid essential ETS family of transcription factors (Doré and Crispino 2011). At the core of this network is the GATA switch, which acts as the fork in the cellular differentiation path to either erythrocyte or megakaryocyte (Doré and Crispino 2011, De Kleer et al. 2014). This GATA switch is intimately related to the ETS family constituents, such as ETS1 and Sp1, as these genes promote megakaryopoiesis at the expense of erythropoiesis (Ikinomi et al. 2000). This ground truth biology suggestive of two pathways based on this switch: a Gata1 enriched pathway with downstream targets such as Klf1 (EKLF), indicative of erythropoiesis, and an ETS-enriched pathway indicative of megakaryopoiesis. The relationship between these cell fate determining pathways being the co-repressive interaction between the GATA and ETS family TFs (Doré et al. 2012).

The third cluster scPectral discovered included Gata1, Klf1, Ets1, Plek, Cebpe, and Gata2. Klf1 and Cebpe are known downstream targets of Gata2, while the Gatas are downstream targets of one another (Doré and Crispino 2011). One of scPectral’s shortcomings is that it cannot describe any nuanced interactions within the pathways it discovers, given that both strong inducer and repressor activity would manifest as high TE scores in locaTE’s network. For this cluster, the known corepressive relationship between the Gatas and ETS suggests that their appearance in the same cluster is due to their strong negative correlation. The constituents of Cluster 3 are however primarily suggestive of the process of red blood cell differentiation. This conclusion is further supported by Metascape’s analysis (Figure 6), with the principal enrichment scores being assigned to erythrocyte and granulocyte differentiation.

**Fig. 6.**
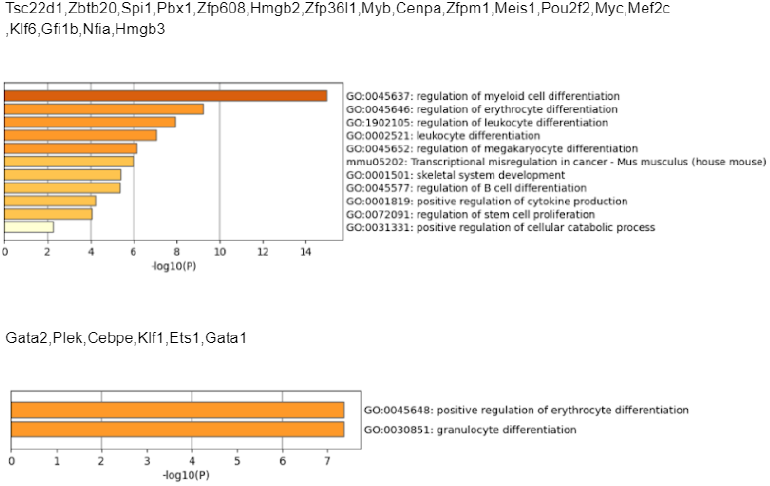
HSPC Metascape

Between the two other clusters, one was not found to be significant in any known pathways, and the other contained contradicting results (Figure 6) such as including genes overpressed in both erythrocyte and megakaryocyte differentiation, two mutually exclusive processes (Doré and Crispino 2011).

## Discussion

It is this second analysis, and particularly its comparison to IQCELL (Heydari et al. 2022), which similarly aims to determine the gene-gene interactions constituting developmental processes, that are exemplary of where scPectral may find its niche. While scPectral did erroneously group genes from different pathways into the same cluster, an issue which may be addressed by using signed weights, and it is a low resolution method that cannot delineate the 16 specific interactions within the erythroid differentiation pathway (Heydari et al. 2022), it did nonetheless show some success at identifying the key constituents of different developmental pathways. Benchmarking scPectral then is difficult, while methods such as IQCELL can report a 68% success rate in discovering known interactions, scPectral can only be validated insofar as it finds the genes involved in these interactions. What this generality does is make scPectral potentially suitable, not as a replacement for existing GRN inference techniques, but rather as a supplementary means by which exploration of novel pathways can begin.

## Conclusion

Presented here has been a novel approach to determining developmental pathways. scPectral falls under the umbrella of GRN inference, and the principal contribution it offers lies in its ability to identify genes involved in developmental cascades while requiring no biologically-guided domain knowledge to steer its analysis. The model achieves this by addressing two frequently neglected aspects of existing GRN inference methods: robust representation of TF interactions and a dynamically-informed metric to weight such interactions (Stumpf 2021). The latter is accounted for by using locaTE’s TE-weighted network as input, and the former through scPectral’s hypergraph abstraction. A hypergraph-adapted spectral clustering algorithm (Hayashi et al. 2020, Zhou and Schölkopf 2007) was used to identify the communities proposed to represent developmental cascades: the GRN motif that is a hallmark of any development process (Alon 2020). Proper validation of these findings requires experimental investigations such as perturbation analysis; however, given the resource constraints, in their stead scPectral was benchmarked by making use of appropriate stem cell differentiation data (Hayashi et al. 2018, Marot-Lassauzaie et al. 2022). With the extensive literature available pertaining to these datasets, scPectral’s efficacy was determined by comparing its clusters to known TF relationships, and by running pathway enrichment analysis using Metascape on these clusters. ScPectral showed mixed results in its ability to determine the developmental relationships contained within the input datasets.

ScPectral’s modest reliability in determining meaningful pathways could potentially be addressed by modifying how it builds hypergraphs to represent GRNs. First, as mentioned previously, high TE scores had no indication of whether the relationship involved activation or repression, and this had the undesirable effect of grouping genes which inhibit one another into the same cluster. Adding signed weights to the TE scores is a potential solution to the issue. The next, and more glaring, issue with the methodology presented lies in its reliance on an initial pairwise network. Generally, pairwise graphs are inferred from hypergraphs, not the other way around (Hayashi et al. 2020). A more suitable approach, making full use of the hypergraph data structure, would be to begin with an initial representation as a hypergraph; specifically, one with both weighted hyperedges and edge-dependent vertex weights (Hayashi et al. 2020). Transfer entropy could be incorporated to the latter to account for gene-specific dynamics, while the former would add a layer to account for the effect of entire TF groups on network behaviour. Another limiting aspect of scPectral’s hypergraph representation is its restriction concerning repeated genes. Every gene could only appear in one pathway, and it could only appear in that pathway once, meaning that a gene could only appear in the output, across all clusters, at most one time. This severely limits the kind of pathways this method can potentially discover. For example, in the case of haematopoiesis, a process which involves a known set of 16 interactions involving seven key genes (Heydari et al. 2022), necessitates that certain genes appear in more than one of these interactions. This specification is one scPectral cannot satisfy. While the purpose of the method presented was to identify developmental cascades, for which scPectral’s behaviour is mostly aligned, future work could look to make the method more flexible, which, with a more robust initial representation, could enable it to identify a wider variety gene network motifs (Alon 2020).

To conclude, the method presented is not a replacement for existing approaches to characterise interaction networks; instead, it has potential as a supplementary tool to aid the process of exploration in determining novel developmental pathways. With appropriate experimental validation, manifest as perturbation experiments, the approach presented may therefore find success as an early step in the process of elucidating novel, as yet characterised, developmental pathways. Future extensions could use the hypergraph more effectively to enable more reliable determination of a wider array of genetic pathways, to better aid experimentalists conducting exploratory investigations.

## Acknowledgment

The author would like to thank Dr. Andrew Roth, in addition to his classmates and colleagues, particularly Kaiyun Guo and Anton Afanassiev, for the beneficial feedback they provided when taking the time to review this work in its early stages.

## Notes

### Competing Interest Statement

The authors have declared no competing interest.

## References

1. Alon U. An Introduction to Systems Biology: Design Principles of Biological Circuits. Boca Raton: CRC Press Taylor amp; Francis Group 2020.

2. Bassalert C, Valverde-Estrella L, Chazaud C. Primitive endoderm differentiation: From specification to epithelialization. Current Topics in Developmental Biology 2018; 128, pp. 81–104.

3. De Kleer I, Willems F, Lambrecht B, and Goriely S. Ontogeny of Myeloid Cells.Frontiers in Immunology 2014.

4. Doré LC, Crispino JD. Transcription factor networks in erythroid cell and megakaryocyte development. Blood 2011; 118 (2): 231–239.

5. Doré LC, Chlon TM, Brown CD, White KP, Crispino JD. Chromatin occupancy analysis reveals genome-wide GATA factor switching during hematopoiesis. Blood 2012; 119 (16): 3724–3733.

6. Gilbert SF. Developmental Biology. 6th edition. Sunderland (MA): Sinauer Associates 2000; Endoderm.

7. Hayashi T, Ozaki H, Sasagawa Y et al. Single-cell full-length total RNA sequencing uncovers dynamics of recursive splicing and enhancer RNAs. Nat Communications 2019; 9, 619.

8. Hayashi K, Aksoy SG, Park CH, Park H. 2020. Hypergraph Random Walks, Laplacians, and Clustering. Proceedings of the 29th ACM International Conference on Information and Knowledge Management (CIKM ‘20). Association for Computing Machinery 2020; New York, NY, USA, 495–504.

9. Hermitte S, Chazaud C. Primitive endoderm differentiation: from specification to epithelium formation. Philos Trans R Soc Lond B Biol Sci. 2014; Dec 5;369(1657):20130537.

10. Heydari T, Langley MA, Fisher CL, Aguilar-Hidalgo D, Shukla S, Yachie-Kinoshita A, Hughes M, McNagny KM, Zandstra PW. IQCELL: A Platform for Predicting the Effect of Gene Perturbations on Developmental Trajectories Using Single-Cell RNA-Seq Data. PLOS Computational Biology 2022; 18, no. 2.

11. Ikonomi P, Rivera CE, Riordan M, Washington G, Schechter AN, Noguchi CT. Overexpression of GATA-2 inhibits erythroid and promotes megakaryocyte differentiation. Exp Hematol, 2000; vol. 28.

12. Kim J, Jakobsen ST, Natarajan KN, Won KJ. TENET: gene network reconstruction using transfer entropy reveals key regulatory factors from single cell transcriptomic data. Nucleic Acids Res. 2021; Jan 11;49(1).

13. Luecken MD, Theis FJ. Current Best Practices in Single-Cell RNA-SEQ Analysis: A Tutorial. Molecular Systems Biology 2019; 15, no. 6.

14. Luxburg U. Tutorial on spectral clustering. Stat. Comput. 2007; 17, 1–32.

15. Marot-Lassauzaie V, Bouman BJ, Donaghy FD, Demerdash Y, Essers MAG, et al. Towards reliable quantification of cell state velocities. PLOS Computational Biology 2022; 18(9):e1010031.

16. Michoel, Tom, and Bruno Nachtergaele. “Alignment and Integration of Complex Networks by Hypergraph-Based Spectral Clustering.” Physical Review 2022; E 86, no. 5.

17. Ng AY, Jordan MI, Weiss Y. On spectral clustering: Analysis and an algorithm. Proc. 15th Annu. Conf. Neural Inf. Process. Syst. 2001; pp. 849–856.

18. Purnick, P, Weiss, R. The second wave of synthetic biology: from modules to systems. Nat Rev Mol Cell Biol 2009; 10, 410–422.

19. Schreiber T. Measuring Information Transfer. Physical Review Letters 2000; 85, no. 2: 461–64.

20. Xiaojie Q, Rahimzamani A, Wang L, Ren B, Mao Q, Durham T, McFaline-Figueroa JL, Saunders L, Trapnell C, Kannan S. Inferring Causal Gene Regulatory Networks from Coupled Single-Cell Expression Dynamics Using Scribe. Cell Systems 2020; 10, no. 3.

21. Stumpf MPH. Inferring Better Gene Regulation Networks from Single-Cell Data. Current Opinion in Systems Biology 2021; 27:100342.

22. Wenjing S, Wang H, Pan G, Geng Y, Guo Y, Pei D. Regulation of the Pluripotency Marker Rex-1 by NANOG and SOX2. Journal of Biological Chemistry 2006; 281, no. 33: 23319–25.

23. Zhang SY, Stumpf MPH. Dynamical Information Enables Inference of Gene Regulation at Single-Cell Scale. bioRxiv 2023; 01.08.523176.

24. Zhang S, Afanassiev A, Greenstreet L, Matsumoto T, Schiebinger G. Optimal transport analysis reveals trajectories in steady-state systems. PLOS Computational Biology 2021; 17(12):e1009466.

25. Zhou D, Huang J, Schölkopf B. Learning with hypergraphs: Clustering, classification, and embedding. Advances in Neural Information Processing Systems 2007; pp. 1601–1608.

26. Zhou Y, Zhou B, Pache L et al. Metascape provides a biologist-oriented resource for the analysis of systems-level datasets. Nature Communications 2019; 10, 1523.

